# Construction of an embryonic caudal organizer by BMP4

**DOI:** 10.1101/2024.11.07.622421

**Authors:** Yan-Yi Xing, Ying Huang, Tao Cheng, Tao Luo, Yang Dong, Zi-Xin Jin, Yi-Meng Tian, Xiang Liu, Jun Ma, Jun-Feng Ji, Peng-Fei Xu

## Abstract

The caudal part of a vertebrate embryo consists of somites, neural tube, lateral plate mesoderm derivatives and the tailbud. However, the minimal conditions and factors sufficient to induce the caudal region, particularly in humans, remain unresolved. Here, we show that BMP4 alone, when administered at appropriate dosage, is sufficient to induce the formation of an organizer for caudal induction. This organizer induces caudal cell fate specification and morphogenesis in zebrafish embryos. In 3D human pluripotent stem cells (hPSCs) aggregates, BMP4 can induce an elongated embryonic structure characterized by caudal fates. Importantly, hPSCs instructed by BMP4 are sufficient to induce a secondary caudal region when grafted into the animal pole of the zebrafish embryo. Our study thus uncovers BMP4 as the inducer in the formation of a caudal organizer in the vertebrate embryo.

**Significance statement:** We demonstrate that BMP4 alone can induce the formation of a caudal organizer, a critical structure that guides the development of the caudal region in vertebrates. Using zebrafish embryos, embryonic explants, human pluripotent stem cell xenografts, and 3D human stem cell aggregates, we show that this organizer replicates morphogenesis and key differentiation pathways seen in natural development. Our findings uncover a novel role for BMP4 and define the minimal requirements for inducing this organizer, offering new insights into vertebrate development and potential applications in regenerative medicine.

## Introduction

During early vertebrate embryogenesis, body axis elongation occurs in two principal stages. Initially, lineage specification during gastrulation results in the formation of three germ layers, where processes like involution, convergence and extension shape and elongate these layers. This leads to the development of the head and trunk regions, with mesodermal precursors contributing to the axial, paraxial, and lateral plate mesoderm (LPM)(1). Subsequently, a structure known as the tailbud emerges at the posterior embryo end, harboring progenitors vital for cell lineage specification and tail elongation, which forms the caudal body axis. In zebrafish, tailbud formation occurs as the blastoderm margin converges and fuses at the end of gastrulation(2, 3). The developmental trajectory of the tailbud and the critical signals regulating its formation remain unclear. Additionally, our understanding of caudal region formation in humans is still very limited.

A century ago, Mangold and Spemann discovered that the dorsal blastopore lip of an amphibian gastrula can induce a secondary axis (mainly the head and trunk region)(4). After that, the organizer concept has been well recognized in different developmental processes and stages, across different species, and the molecular natures of those organizers have also been well characterized(5, 6). However, the minimal conditions that can induce an organizer capable of inducing and patterning the caudal part of the vertebrate embryo have not been addressed. Stem cell-based synthetic embryology has created new research models and provided significant insights into developmental and regenerative biology(7). By applying essential morphogens globally or locally, key aspects of early development have recently been recapitulated *in vitro* with pluripotent stem cells. For instance, activation of the WNT signaling pathway by its agonist treated homogenously can result in the formation of gastruloids(8), trunk-like structures(9), or somitoids(10), depends on the culture conditions. Similarly, the development of A-P patterned head-like structures can be induced by a single gradient of Nodal(11). Treatment with BMP4 in micropatterned human stem cells can recapitulate early embryonic spatial patterning(12) and BMP4 induced morphogen signaling center could result to the development of a mammalian embryo model(13). Moreover, human stem cell colonies treated with Wnt and Activin can function as ‘organizer’(14).These models have effectively mimicked germ layer specification, body patterning and somitogenesis.

BMP4, a member of the TGF-β superfamily, has been shown to play critical roles in body plan establishment and tailbud development across various organisms such as mice(15), frog(16), chick(17), and zebrafish(18). In this study, we demonstrate that a moderate BMP4 signaling can induce the formation of a secondary axis in zebrafish that contains caudal structures yet lacks axial mesoderm and endoderm. Similar results can be obtained from the BMP4-induced zebrafish explant and 3D aggregates of human pluripotent stem cells. By using single-cell RNAseq and live imaging, we systematically analyzed the dynamics of cell fate specification, cell movements of key cell types during those processes. Additionally, the organizing potential of BMP4-treated hPSCs was further validated through xenograft experiments.

## Results

### Bmp4 induces a secondary tail in zebrafish embryos

We firstly employed zebrafish animal pole blastomeres as a model to explore the organizing capacity of Bmp4. The pluripotent, naïve, and equivalent nature of the animal pole cells in zebrafish embryos makes them an ideal system for studying germ layer induction and morphogenesis(19).

By injecting 10 pg of zebrafish *bmp4* mRNA into one animal-pole blastomere at 128-cell stage, we established a BMP signaling gradient (Figure S2A). No obvious morphological changes were observed in injected embryos from the blastula stage until the beginning of gastrulation (6 hpf, hours post fertilization). Interestingly, a protrusion progressively bulges at the injection site and becomes evident at mid-gastrulation (8 hpf, Figure 1A). At the beginning of the segmentation stage (12 hpf), a distinct secondary axis became apparent (Figure 1A). This axis extends from the head region of the primary axis to a separate protrusive end that resembles the tailbud. By 24 hpf, the *bmp4*-injected embryo developed a secondary axis that diverged from the main axis at the head region, featuring multiple somites but lacking a visible notochord, indicating the absence of axial mesoderm (Figures 1A and S1B).

**Figure 1.**
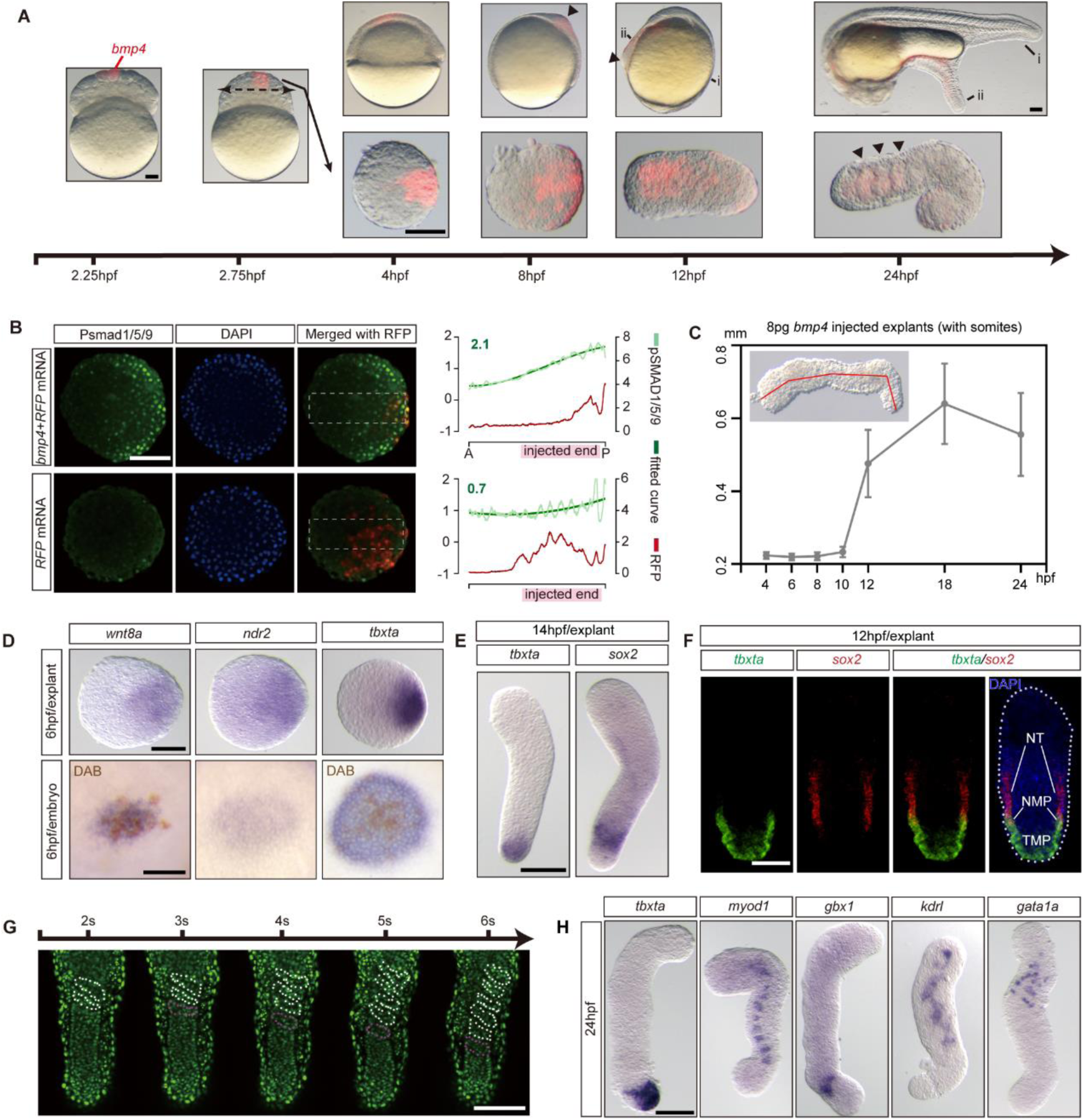
Testing induction capability of BMP4 in zebrafish embryo and explant. (A) Representative images of bmp4 injected zebrafish embryos (upper) and explants (lower, with injection site towards the right) from 2.25hpf to 24hpf. Red fluorescence marks the descendants of the bmp4 injected blastomere. Arrowheads at 8hpf and 10hpf embryo point to the protrusions induced by BMP4 in embryos, and at 24hpf in explants to the segmented somites. Lables indicate the primary (i) and secondary axis (ii). (B) Immunofluorescence imaging of Phospho-smad1/5/9 in *bmp4+RFP* injected explant (upper) and *RFP* injected explant (lower) at 4 hpf, with injection site towards the right. Signal measurements along the doted rectangles are shown on the right. Dark Green curves represent the nonlinear fit of actual data curves (light green), with Dark Green numbers indicating the mean slopes. Red curves depict the distributions of RFP signaling intensity along the doted rectangles. (C) Quantification of the elongation of bmp4 explants from 4hpf to 24hpf. The Midlines along anterior posterior axis of bmp4 explants are measured (red line), as displayed in the upper left. (D) bmp4 injected explants (upper, posterior towards the right) and embryos (lower, animal pole view) at 6hpf stained by whole mount *in situ* hybridization (WISH) with the indicated probes. DAB staining of GFP (brown spots) indicates the clones of *bmp4*&*GFP* mRNA injected blastomere at 2.25 hpf. (E) bmp4 explants at 14 hpf stained by WISH with indicated probes. (F) Whole mount *in situ* hybridization chain reaction (HCR) staining of bmp4 explants at 12hpf for *tbxta* and *sox2*, with co-staining of DAPI. NT, neural tube; MP, neuromesodermal progenitor; TMP, tailbud mesodermal progenitor. (G) Time-lapse imaging of bmp4 explants displaying the sequential formation of somites, as per supplementary move S11. White doted lines indicate existing somites, and purple doted lines indicate newly formed somites. (H) bmp4 explants at 24 hpf stained by WISH with the indicated probes. For all explants displayed in this article, anterior (opposite to the injection site) is towards the top unless otherwise stated. Scale bars: 100 μm.

Molecular analysis of the induction process revealed that Bmp4 activates key morphogenetic signals pivotal for early development, including Nodal, Wnt and FGF, as evidenced by the expression of *ndr2*, *wnt8a* and *fgf8a* (Figures 1D and S2D). At the same time, the naive animal pole cells are transformed into ventral mesoderm in response to Bmp4, as shown by the expression of *eve1* (Figure S2D). However, we failed to detect the expression of endoderm marker *sox17* near the *bmp4* injected region (Figure S2D).

A secondary tailbud induced by Bmp4 is evident at 12 hpf, marked by the expression of *tbxta* (Figure S2E). At 24 hpf, markers of ventral-lateral mesoderm derivatives such as pronephric duct (*cldn3d,* Figure S2I) and somites (*tnnt2c*, *myod1,* Figure S2H) can be detected in the secondary axis. However, specification of axial mesoderm is not observed, as indicated by the absence of *shha* expression (Figure S2G). Additionally, a continuously dorsalized neural tube extending to the tail tip is observed, marked by the expression of *sox19a* and *olig2* (Figure S2F). Notably, no specification of a head region is detected in the secondary axis, as indicated by the lack of *egr2b* expression (Figure S2F).

### *in vitro* induction of a caudal-like structure by moderate Bmp4 signaling

To further explore the organizing potential of Bmp4 *in vitro*, we explanted the animal pole cells from embryos injected with *bmp4* mRNA at 512-cell stage (hereafter referred to as ‘Bmp4 explant’). We then investigated the morphological and cellular events within these explants.

Upon excision, the explant rapidly became a spherical shape, and established a BMP signaling gradient (Figures 1B and S2B). Begin from the onset of segmentation, the explants exposed to a moderate amount (8-10 pg of mRNA) of *bmp4* exhibited a remarkable elongation (about 50% of injected explants, Figure S1F and supplementary movie 1), whereas both insufficient and excessive levels of *bmp4* failed to stimulate this elongation. (Figures 1A, 1C, S1A, S1C-M). Interestingly, the elongated explants exhibited sequential somite formation (Figure 1G and supplementary movie 11).

Consistent with the *in vivo* results, the expression of *wnt8a*, *ndr2* and *eve1* in the Bmp4 explants was also activated during gastrulation (Figure 1D). At the segmentation stages, we found that *tbxta* was expressed at one end of Bmp4 explant, which we designated as the posterior. Additionally, A neural domain expressing *sox2* was found to partially overlap with a tail bud mesodermal domain expressing *tbxta,* but expressed slightly more anteriorly, suggesting the induction of neuro-mesodermal progenitors (NMP, *tbxta*^+^&*sox2*^+^, Figures 1E and 1F).

By 24 hpf, we detected the specification of the ventral-lateral mesodermal derivatives such as erythrocyte (*gata1a*) and vascular endothelium (*kdrl*). And the Bmp4 explants eventually formed 8-10 somites, as indicated by *myod1* expression (Figures 1H, S2O). However, we did not detect the formation of notochord or head region (*tbxta*, *egr2b*, Figures 1H, S2L). Although a neural domain expressing *gbx1* was found in the explant, it was not well organized compared to the *in vivo* neural tube (Figure 1H). Taken together, these results indicate that Bmp4 explant closely resemble the caudal region of the zebrafish embryo.

### Emergence of the anterior-posterior axis of Bmp4 explant

To systematically analyze the cell fate specification of Bmp4 explant, we performed single cell sequencing analysis. We collected 35,830 single-cell transcriptomes from 6 developmental stages of Bmp4 explants over 24 hours (Figures 2A, S3A, supplementary table 1), and 28 cell clusters were identified and annotated based on the marker genes expression. These clusters could be categorized into somitogenesis-related clusters, lateral plate mesoderm and its derivatives, neural ectoderm and its derivatives, epidermal ectoderm and its derivatives, and other cell clusters (Figure 2A).

**Figure 2.**
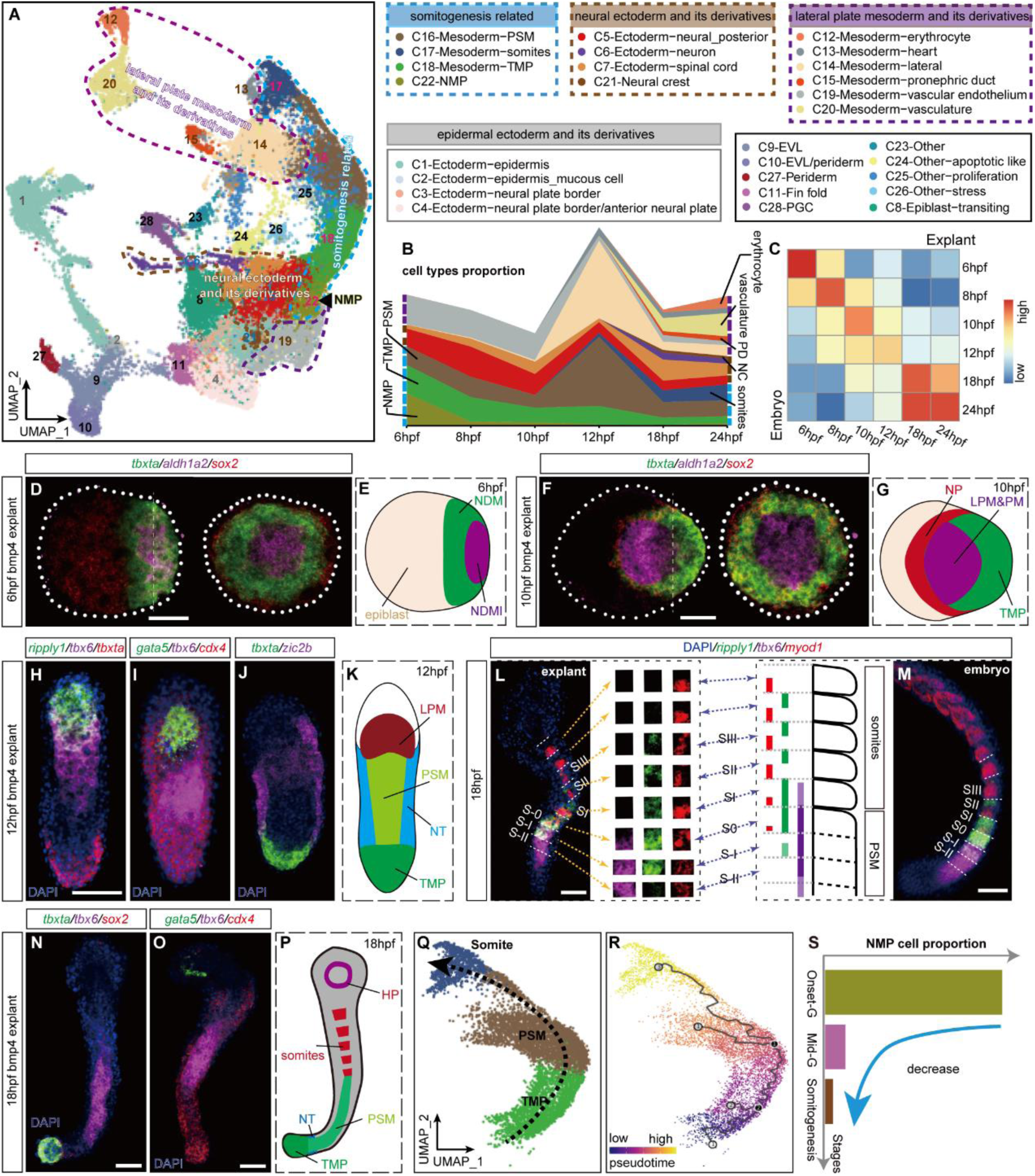
Analysis of cellular composition and pattern formation dynamics in Bmp4 explants. (A) Uniform manifold approximation and projection (UMAP) of integrated scRNA-seq datasets for Bmp4 explants at 6hpf, 8hpf, 10hpf, 12hpf, 18hpf and 24hpf, colored by cell clusters. Blue, brown and purple dotted circles indicate clusters related to somitogenesis, neural ectoderm/ derivatives, and lateral plate mesoderm/derivatives, respectively. Arrowhead marks the neural mesodermal progenitors (NMP). (B) Ribbon plot showing the temporal proportions of cell clusters related to somitogenesis and lateral plate mesoderm/derivatives in Bmp4 explants. (C) Heatmap showing the Pearson correlation between zebrafish embryos and Bmp4 explants at indicated developmental stages, showcasing the similarity in gene expression patterns. (D, F) Left: HCR co-staining of Bmp4 explants at 6hpf (D) and 10hpf (F) for *tbxta*, *aldh1a2* and *sox2*, with posterior to the right. Right: Sagittal views (from the posterior end) of the same stained explant in the left panel along the dashed lines. Dashed circles demarcate the boundaries of explants. (H-J) HCR staining for *ripply1*, *tbx6* and *tbxta* (H), *gata5*, *tbx6* and *cdx4* (I), *tbxta* and *zic2b* (J) of Bmp4 explants at 12hpf. (L, M) HCR staining of Bmp4 explant (L) or zebrafish embryo (M) at 18hpf for *ripply1*, *tbx6* and *myod1*, oriented with anterior towards the top. Somites are segmented by white dashed lines. Expression patterns of 3 genes in each somite of Bmp4 explant are displayed as insets (L, right) and schematically for the embryo (M, left). (N, O) HCR co-staining for *tbxta*, *tbx6* and *sox2* (N), and *gata5*, *tbx6* and *cdx4* (O) in Bmp4 explants at 18hpf. (E, G, K and P) Schematic summaries illustrating patterns of cell clusters in Bmp4 explants at 6hpf (E), 10hpf (G), 12hpf (K), 18hpf (P). NDM, non-dorsal margin; NDMI, non-dorsal margin involuted; LPM, lateral plate mesoderm; PM, paraxial mesoderm; PSM, pre-somitic mesoderm; HP, heart progenitor. All samples in (H-J, L-O) were co-stained with DAPI. (Q, R) Cell state trajectory analysis of somitogenesis-related cell clusters. Cells are colored by cell types (Q) and pseudotime (R). (S) Time-course analysis of the proportion of NMP cells in bmp4 explants. Scale bars:50 μm.

It is noteworthy that lineages such as paraxial mesoderm related progenitors, including NMP, tailbud mesoderm progenitor (TMP(20)) and pre-somitic mesoderm (PSM) were specified during gastrulation stage, and gradually diminished during somitogenesis stage. In contrast, tissues such as somites, pronephric duct (PD), vasculature and erythrocyte emerged and expended during the organogenesis stage (Figure 2B), which is in accordance with the *in vivo* situation. Bmp4 explant showed transcriptional similarity to wild-type embryos at each of the six developmental time points (Figure 2C), suggesting a comparable developmental trajectory up to 24 hpf. And the developmental time points of Bmp4 explant can be divided into three major stages based on transcriptional similarity: onset-Gastrulation (6 hpf), Mid-Gastrulation (8-10 hpf), and Somitogenesis (12-24 hpf) (Figure S4B).

To elucidate the spatial organization of cell types and their developmental trajectories, we performed hybridization chain reaction (HCR) analysis of key cell clusters at 6 developmental stages. The Bmp4 explant exhibits anterior-posterior (AP) polarity at the onset of gastrulation, with a distinct mesodermal region assigned to the posterior end, which could be further divided into a central domain expressing *aldh1a2* and a surrounding domain expressing *tbxta* (Figures 2D, 2E). By the end of gastrulation (10 hpf), the *aldh1a2*-expressing cell cluster has shifted completely anterior to the *tbxta*-expressing domain, resulting to the formation of anterior-posterior axis of Bmp4 explant (Figures 2F, 2G). Single-cell trajectory analysis indicates that the *aldh1a2*-expressing cell cluster exhibits signatures of paraxial and lateral plate mesoderm (Figure S3H). Additionally, the neural progenitors, specified during the gastrulation stage, flanks the entire mesodermal region (Figures 2F, 2G). It is noteworthy that the mesodermal and neural domains specified in Bmp4 explant are radially symmetrical viewed from the sagittal section, suggesting the lack of left-right axis (Figures 2F, 2G).

### Recapitulation of somitogenesis in Bmp4 explant

Somitogenesis is a highly conserved developmental process across vertebrates, characterized by a sequential progression of differentiation from head to tail (21). During this process, new somites bud off from the anterior part of the presomitic mesoderm (PSM), while a continuous influx of cells from the tailbud replenishes and sustains to the posterior PSM. These formed somites then differentiate into various tissues, including vertebrae, muscles, and dermis at later stages of development (22).

In Bmp4 explant at the early segmentation stage (12-14 hpf), a distinct PSM region (marked by *tbx6* and *her1)* can be observed located posterior to the LPM (marked by *gata5*) and anterior to the TPM (*tbxta*^+^*sox2*^-^*zic2b*^-^) (Figures 2H-K, S4F). By 18 hpf, Bmp4 explant exhibited distinctive molecular characteristics that made it easily distinguishable between mature, newly formed, and prospective somites(23). *tbx6* is highly expressed throughout the anterior PSM (S0, -I, -II), decreases in the newly formed somite (SI), and becomes undetectable beyond SII. The expression of *ripply1* initiates at the posterior of S-I, upregulated throughout S0 and S1, restricted to the anterior half of SII, SIII, and becomes barely detectable beyond SIII. In most cases, *myod1* expression is restricted to the posterior half of the somite in Bmp4 explant, indicating the onset of myogenesis and the proper establishment of anterior-posterior polarity in the somites (Figures 2L, 2M). In addition, *cdx4* expression extends from the TPM to the *gata5*^+^ domain, suggesting that the majority of Bmp4 explant exhibits ventral and posterior identities (Figures 2N-P).

Single-cell trajectory analysis also revealed a clear differentiation trajectory that initiates from tailbud cell state, progresses though PSM, and ultimately ends at somites state (Figure 2Q,2R). Along this trajectory, we identified many genes that remain poorly studied regarding their roles in somitogenesis, in addition to those genes previously reported to play roles in the differentiation from TPM to PSM and from PSM to somites. Thus, Bmp4 explant hold great potential as a valuable model for future research in somitogenesis (Figure S3C-G, S3I-K).

NMP cells in zebrafish have been demonstrated to be bipotential. However, One NMP cell rarely give rise to both neural and mesodermal decedents, due to its low proliferation frequency (24). We also identified the presence of NMP cells expressing *tbxta* and *sox2* in Bmp4 explant and observed similar biological characteristics of NMP cells in zebrafish (Figures 1F, S4A, S4C, S4D). The proportion of NMP cells decreased quickly from gastrulation to segmentation stages (Figures 2N,2P,2S,3G, S3B) and they tended to stay in G1 phase rather than advancing to S phase, in contrast to TMP and neural cells (Figure S4E), suggesting the low proliferating frequency of the NMP cells.

**Figure 3.**
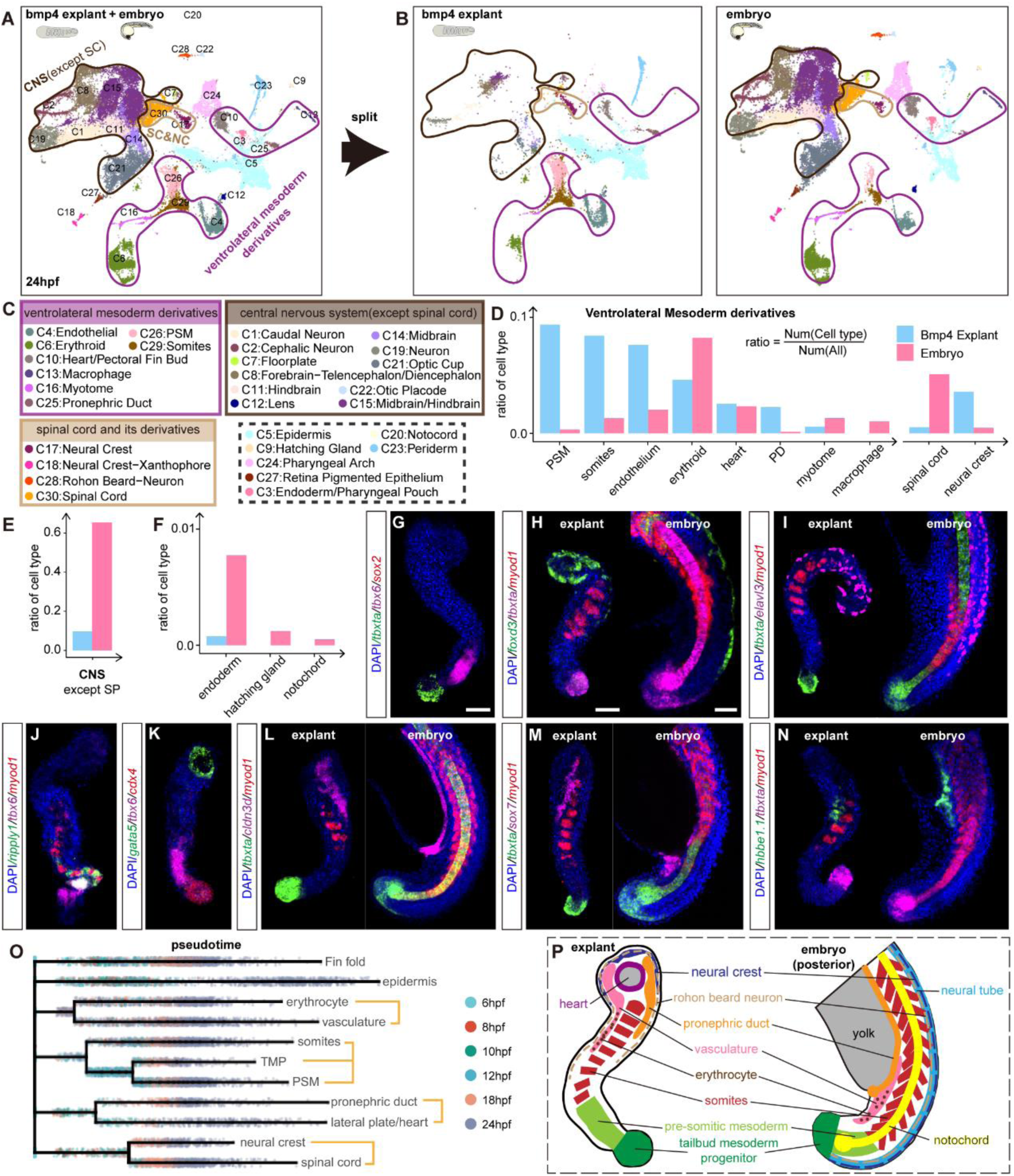
Characterization of Bmp4 explants at the organogenesis stage. (A, B) UMAP plots showing integrated single-cell RNA sequencing (scRNA-seq) datasets of zebrafish embryos and Bmp4 explants at 24 hpf. (A) Displays the combined datasets with cells colored by the cell clusters listed in (C). (B) Shows separated datasets for Bmp4 explants (left) and zebrafish embryos (right). Black, purple, and yellow circles highlight cell clusters of the central nervous system (excluding spinal cord), ventrolateral mesoderm derivatives, and spinal cord & its derivatives, respectively. (D-F) Comparisons of the proportions of indicated cell clusters between Bmp4 explants and zebrafish embryos at 24hpf. PD, pronephric duct; SP, spinal cord; NC, neural crest; CNS, central nervous system. (G, J, K) HCR co-staining for *tbxta*, *tbx6* and *sox2* (G), *ripply1*, *tbx6* and *myod1* (J), *gata5*, *tbx6* and *cdx4* (K) in bmp4 explants at 24hpf. (H, I, L-N) HCR co-staining for *foxd3*, *tbxta*, and *myod1*(H), *tbxta*, *elavl3* and *myod1*(I), *tbxta*, *cldn3d* and *myod1*(L), *tbxta*, *sox7* and *myod1*(M), *hbbe1.1*, *tbxta* and *myod1*(N) in Bmp4 explants (left) at 24hpf and zebrafish embryos (right, only the posterior part is displayed) at 20-24hpf. Anterior is towards the top for both the explants and embryos. Samples in (G-N) were all co-stained with DAPI. (O) Force-directed layout of Bmp4 explant scRNA-seq data. Cells are color coded by developmental stages, with terminal cell clusters labeled by cell identities. (P) Schematic summary illustrating structures of Bmp4 explant at 24hpf, compared to caudal part of zebrafish embryo. Scale bars:50 μm.

Collectively, these findings demonstrate that Bmp4 explant exhibit morphological and molecular characteristics of the posterior embryo and undergo a somitogenesis progression akin to that observed *in vivo*.

### Caudal organ primordia are orderly organized in Bmp4 explant at 24 hpf

By the end of 24 hpf, the zebrafish body plan is well established, with clearly apparent organ primordia (25). To systematically compare Bmp4 explant with the embryo at this stage, we integrated our single-cell datasets from Bmp4 explant with a published embryonic single-cell dataset at 24 hpf (Figures 3A-3C). Unsupervised clustering revealed 30 distinct cell clusters. We observed that the majority of cells in Bmp4 explant are ventrolateral mesoderm derivatives, including somites, endothelium, pronephric duct, erythroid and other related cell types (Figure 3D). In contrast, brain cells (forebrain, midbrain, hindbrain, etc.) were minimally present in Bmp4 explant, with only a few posterior neural cells (such as the spinal cord and neural crest) being detectable. (Figures 3D,3E). Notably, no endoderm or axial mesoderm derivatives (pharyngeal pouch, hatching gland and notochord) were detected in Bmp4 explant (Figure 3F).

To determine whether the organ primordia in Bmp4 explant share similar developmental trajectories with their *in vivo* counterparts, we employed two computational methods to reconstruct the single-cell developmental trajectories of Bmp4 explant. First, we utilized a simulated diffusion-based computational approach (URD), which traced the origins of the tailbud mesoderm progenitor (TMP), presomitic mesoderm (PSM), and somites back to a shared cluster of progenitors at an earlier stage. Similarly, vasculature and erythrocytes, pronephric duct (PD), heart, spinal cord, and neural crest were observed to originate from distinct progenitor clusters (Figure 3O). Subsequently, we employed K-Nearest Neighbor algorithm (KNN) on our time-series single-cell datasets of Bmp4 explant (See Methods). During the segmentation stage, we observed progressive differentiation of the neural plate into neural crest cells and Rohon beard neurons (12-24 hpf). Notably, the NMP cells, which have the potential to generate both mesoderm and neural ectoderm cells, have already disappeared by the onset of segmentation. This observation may explain the diminishment of the spinal cord during this stage. In addition, a clear progression was observed as the paraxial mesoderm differentiates into PSM, which further differentiates into somites. Furthermore, the TMP contributed concurrently to the PSM during this period (Figure S3B). These results indicate that both trunk and tail somites are specified in Bmp4 explant, with trunk somites originating from the paraxial mesoderm formed during gastrulation, while tail somites are specified from cells in the TMP (26–28).

To investigate the arrangement of the organ primordia within Bmp4 explant, we performed HCR co-staining for key makers of each organ primordium. We observed that cell types related to somitogenesis display a pronounced spatial distribution along anterior-posterior (AP) axis: the tailbud mesoderm progenitor is located at the most posterior end of the Bmp4 explant, while the PSM is positioned anterior to the TMP yet posterior to the somites (Figures 3G,3J), mirroring patterns observed in the embryo. Furthermore, ventrolateral mesoderm derivatives are located at the anterior part of Bmp4 explant: the heart primordium is found at the very anterior end; pronephric duct, vascular endothelium and erythrocytes are located between the somites and the heart region (Figures 3K-N). Pronephric duct is in the trunk region of the embryo. It is noteworthy that the posterior part of the PD overlaps with 2-3 anterior somites in Bmp4 explant, which are thus morphologically identified as trunk somites (Figure 3L). The spinal cord, which is located anterior to the TMP, is barely detectable, but its related derivatives such as neural crest and Rohon-beard neuron, are found at the anterior of the Bmp4 explant, surrounding the ventrolateral mesoderm derivatives (Figures 3G-3I). The normal anterior limits of expression of central/posterior *hox* genes (*hoxd10a/hoxd12a*, Figure S4G) and central/anterior *hox* genes (*hoxc6b/hoxb3a*, Figure S4H) further demonstrate that Bmp4 explant is well organized anterior-posteriorly(29).

In the zebrafish embryo, the anatomical trunk - comprising the heart, pronephric duct, and trunk somites - extends from the posterior of head to the anus, while the tail, which includes tissues such as tailbud, tail somites and vasculature, is located posterior to the anal opening. Comparison of Bmp4 explant with the embryo reveals that the majority of tissues in Bmp4 explant—TMP, PSM, posterior somites, vasculature and erythrocyte—compose a tail-like structure. Other tissues, such as pronephric duct, anterior trunk somites, and spinal cord derivatives, are located near the tail in both Bmp4 explant and embryo. Only the heart is an exclusively anterior structure in the Bmp4 explant (Figure 3P).

Taken together, these results demonstrate that Bmp4 explant exhibits structural and molecular features similar to those of the vertebrate caudal region, and thus we designate our model system as a caudal-like structure (CLS).

### Recapitulation of gastrulation and tail elongation cell movements in BMP4 induced CLS

During embryonic development, coordinated cell movements coupled with cell fate specification dynamically sculpt the evolving morphology of the embryo(2) (Figure 4A).To evaluate how cell movements shape this caudal-like structure and to what extent it recapitulates embryonic development, we conducted timelapse confocal imaging and analyzed cell movements using Imaris. Morphogenesis of Bmp4 explant can be divided into two consecutive phases, gastrulation and segmentation stages (Figures 4D,4L), consistent with *in vivo* development. During the gastrulation stage, cells near the injection site converged toward the anterior of the explant and involuted outside-in very quickly (Figure 4B, supplementary movies S2, S3), leading to the formation of a transient hole reminiscent of the blastopore. This hole, eventually closed near the injection site where the TMP cell cluster resides, reminiscent of the fusion of blastoderm margin *in vivo* (Figures 4C, 2G). Similar processes can also be observed on the animal pole of embryo that injected with *bmp4* (Figures S5A-D, supplementary movies S4, S5).

**Figure 4.**
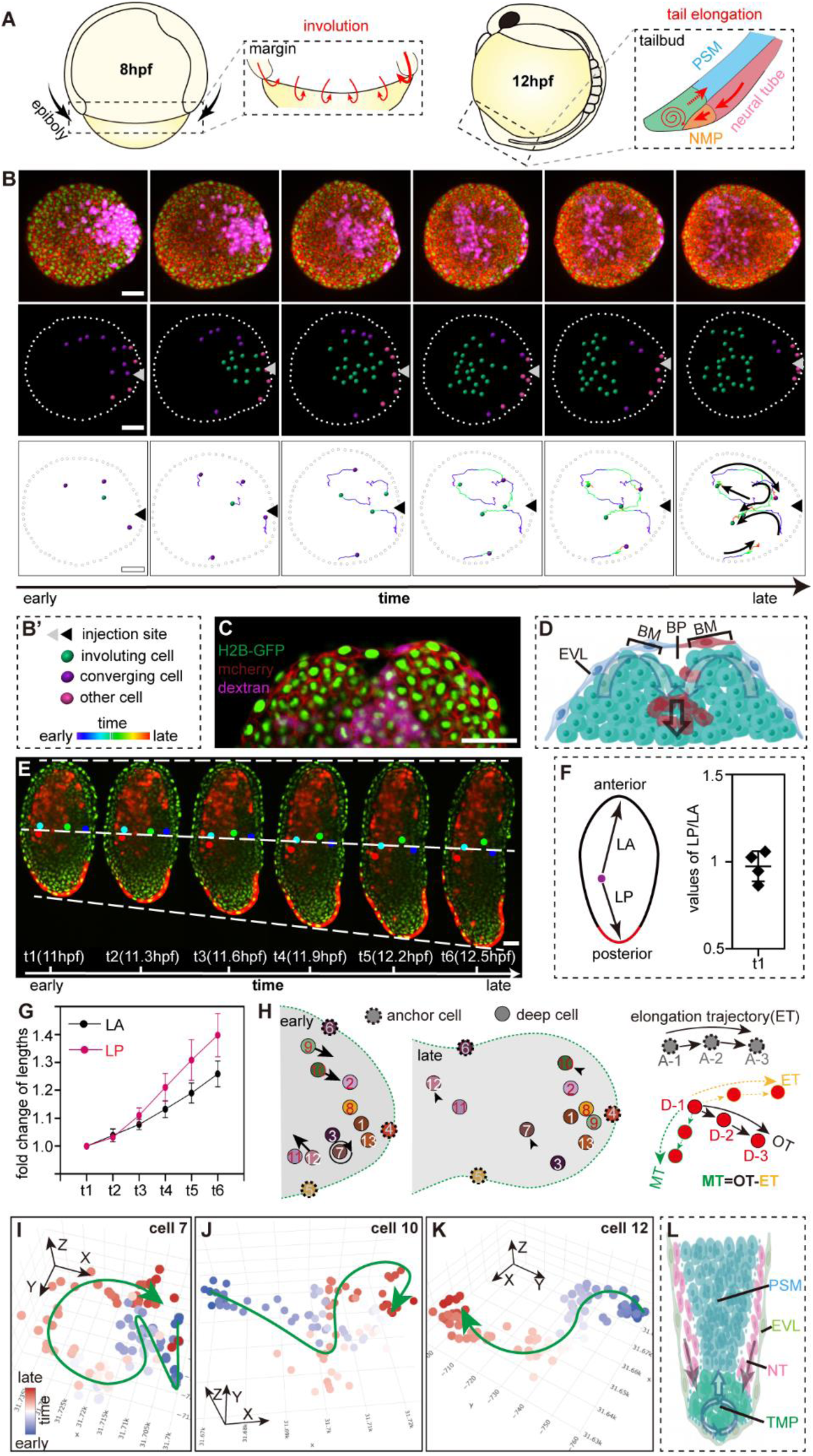
Morphogenetic dynamics of Bmp4 explants. (A) Schematic representation of gastrulation and tail elongation cell movements in zebrafish embryo. PSM, pre-somitic mesoderm. (B) Time-lapse imaging of Bmp4 explants during gastrulation, as detailed in supplementary movies S2 and S3. Upper panel: Representative images of Bmp4 explants with cell nuclei, cell membrane, and descendants of *bmp4*-injected blastomeres labeled with H2B-GFP, mCherry, and Dextran Fluorescein (purple) respectively. Middle panel: Images with selected cells colored by states of cell movements as displayed in panel (B’). Lower panel: Images showing movement trajectories with only five representative trajectories displayed and summarized by white arrows for clarity. The time course of trajectories are color-coded from blue to red as displayed in panel (B’). (C) Representative image of Bmp4 explants during gastrulation, referencing supplementary movie S2. Labeling details are as described in (B). (D) Schematic representation of gastrulation cell movements in Bmp4 explants. EVL, enveloping layer; BM, blastoderm margin; BP, blastopore. Black hollow arrow indicates the direction of involution movement. (E) Time-lapse imaging of Bmp4 explants during segmentation stage. Cell nuclei and descendants of *bmp4*-injected blastomere are labeled with H2B-GFP and dextran fluorescein (red), respectively. Colored spots in the middle panel indicate tracked cells, as shown in supplementary movie S9. White dash lines delineate the most anterior (upper), intermediate (middle), and most posterior (lower) sites of the Bmp4 explant. (F) Measurement of distances from one spot to the anterior or posterior end of the explant, denoted as LA or PA respectively. It is noteworthy that the blastomere injected with *bmp4* mRNA contributes both to EVL cells that stay at the injection site and deep cells that migrate to the anterior later. So, we use the red fluorescence labeled EVL cells to determine the injection site and posterior end. The values of LP divided by LA are all around 1, indicating central positioning of the tracked spots. (G) Quantification of the elongation of the anterior and posterior halves of bmp4 explant over time, with lengths of LP/LA at times t2-6 normalized to those at t1. (H) Schematic representation of positions of tracked cells at early (left) and late (middle) time points. Dashed circles indicate the EVL cells used as anchor cells. Solid circles indicate deep cells. The strategy for analyzing migration trajectories is outlined to the right, including calculation of the elongation trajectory using three anchor cells and subsequent adjustment using the observed trajectories of deep cells to obtain migration trajectories (see Methods). A, anchor cell; D, deep cell; ET, elongation trajectory; OT, observed trajectory; MT, migration trajectory. (H-J) Movement trajectories of cell 7,10,12 respectively, with dots representing the 3D positions of the tracked cells and green lines with arrows manually delineated to indicate the movement trajectories. (K) Schematic of tail elongation cell movements in bmp4 explants. Scale bars: 50 μm.

To investigate which cluster of cells involuted, we tracked the cell movements of a Bmp4 explant constructed from a transgenic line Tg (*aldh1a2*: H2B-RFP). Cells near the injection site exhibited red fluorescence before the onset of involution, suggesting that cell fate specification preceded cell movement for this cluster (Figures S5E, S5F, supplementary movies S6, S7). During gastrulation, *aldh1a2*-expressing cells (LPM and PM) progressively involuted, resulting in the formation of a red fluorescent region anterior to the non-involuting, non-red fluorescent TMP region by the end of gastrulation (Figure S5G).

Then, to ascertain the primary driver of axis elongation during the segmentation stage, we first compared the elongation rates of the anterior and posterior parts of the Bmp4 explant respectively. We found that the posterior part elongated much faster than the anterior part (Figures 4E-G), suggesting that the TMP which located at the posterior end contribute significantly to the elongation of the entire Bmp4 explant during segmentation stage. During embryonic tail elongation, cells in the tailbud exhibit local rotational yet forward movements, contributing to the posterior of PSM, and thereby driving the axis elongation(30, 31). We analyzed trajectories of 13 different cells in posterior end of Bmp4 explant, which exhibited different kinds of movement trajectories: EVL cells (cell 4,5,6) and deep cells (cell 1,2,3,7-13) within or near the TMP (Figure 4H). Given that EVL cells are interconnected by tight junctions and typically do not migrate, the consistent, straightforward trajectories observed for all EVL cells (observed trajectories, OT; Figures S5H, S5I; Supplementary Movie S10) reflect overall explant elongation. Thus, the three EVL cells were defined as anchor cells, with their trajectories representing the explant elongation, which is referred to as the “elongation trajectory (ET)”. Subsequently, we calculated the ‘migration trajectories (MT)’ for other cells by subtracting the ET from OT (Figure 5H). It turned out that most cells remained in the TMP during elongation and exhibited stereotypical rotating trajectories (cell 1,2,3,7,8,11 and Figures 4I, S5J), resembling the local rotational cell movements observed in the embryonic tailbud *in vivo*(30). Some cells in the TMP migrated anteriorly in a zigzag pattern and integrated into the PSM (cell 12 and Figures 4K, S5K). Some cells adjacent to the TMP migrated posteriorly into the TMP (cell 9,10 and Figures 4J, S5L).

**Figure 5.**
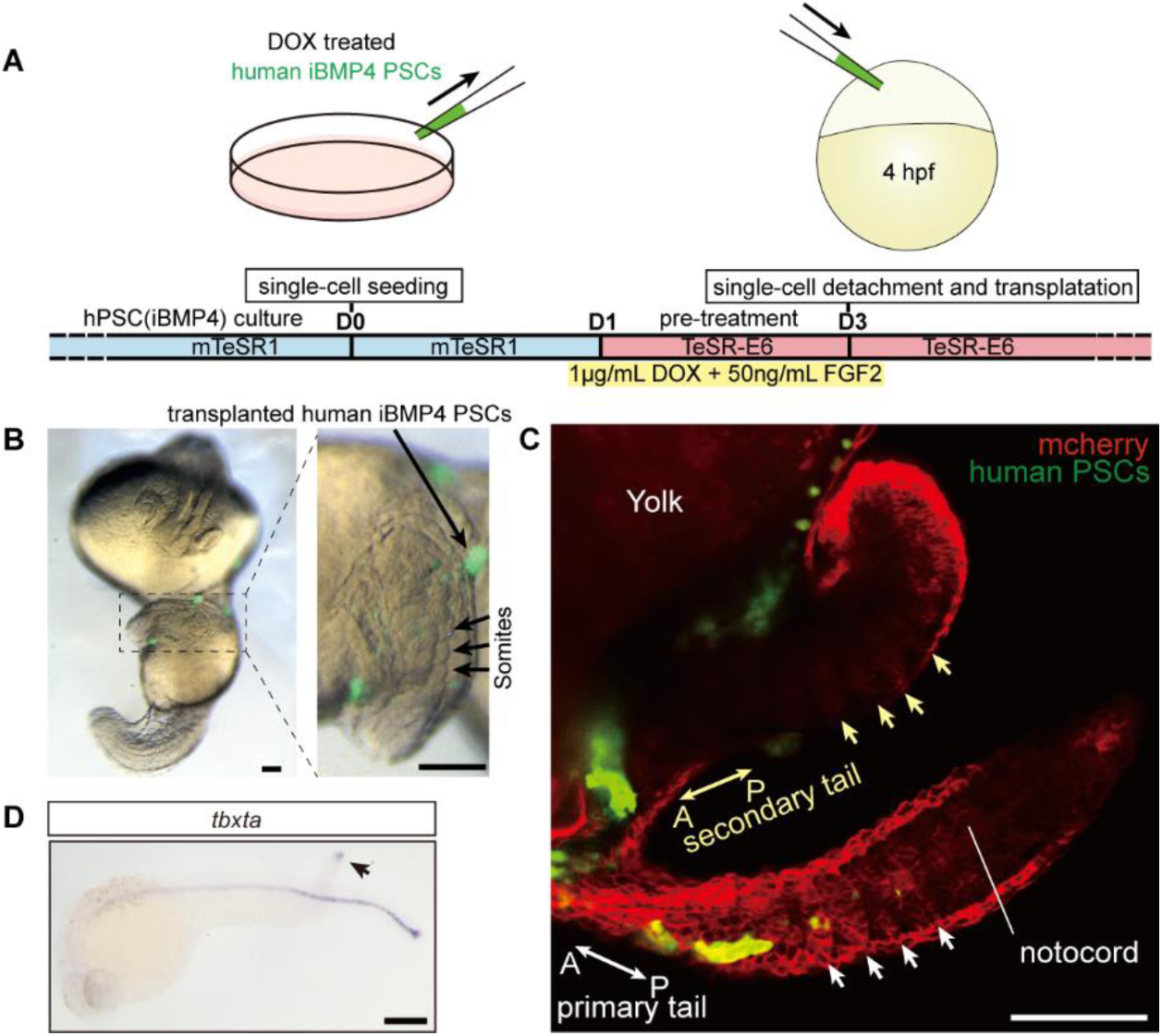
Xenografting BMP4-Treated Human PSCs into Zebrafish Embryos. (A) Schematic illustrating the xenograft procedure of BMP4 treated hPSC into zebrafish embryos. (B) Bright field images showing the secondary tail induced by BMP4 treated hPSCs labeled with GFP. Long black arrow indicates hPSC clones. Short black arrows highlight the somites in the secondary tail. (C) Confocal microscopy image of a zebrafish showing both the primary and secondary tails post-xenograft. Host zebrafish cells and transplanted BMP4-treated hPSCs are labeled with mCherry and GFP, respectively. Arrows indicate the A-P polarity and the somites in the secondary and primary tails, respectively. Notochord was not detected in the secondary axis. (D) WISH of *tbxta* in the xenografted embryo at 24hpf. Arrowhead indicates the tail organizer in the secondary tail. Scale bars:100 μm.

Taken together, these findings suggest that the Bmp4 explant recapitulates not only the structural and molecular characteristics of caudal structures but also the morphogenetic progresses of gastrulation and tail elongation observed *in vivo* (Figures 4A, 4D, 4L).

### BMP4 treated human pluripotent stem cells function as a ‘caudal organizer’

The term ‘organizer’ is functionally defined as a cluster of cells capable of inducing a secondary axis when transplanted ectopically into host embryos (32). The cross-species transplantation strategy is generally accepted as the most stringent test for functionally defining an organizer(14, 33, 34). The induction of a caudal-like structure in the zebrafish explant by Bmp4 prompts us to hypothesize that Bmp4 may induce a ‘caudal organizer’ capable of organizing the formation of caudal structures. To validate this hypothesis and, furthermore, to test whether this function of BMP4 is conserved among vertebrates, we grafted BMP4 treated human PSCs (See Methods) into zebrafish animal poles at sphere stage (Figure 5A). At 24 hpf, we observed that transplanted human cells induced and contributed directly to a secondary tail with sequential somites (Figures 5B, 5C). Whole mount *in situ* hybridization of *tbxta* revealed that TMP is present at the posterior end, and no axial structures are observed in the secondary tail (Figure 5D). Taken together, this functional test demonstrated that BMP4 treated human PSCs meet the criteria of ‘caudal organizer’, and this function of BMP4 is conserved between zebrafish and humans.

### BMP4 induces caudal cell fates and elongation in human 3D PSC aggregates

To further investigate the organizing capability of BMP4 treated human PSCs, we conjugated an aggregation of BMP4-inducible PSCs with an aggregation of normal human PSCs. They fused together quickly and formed an embryoid body (EB) at day 2 (Figure 6A). Under DOX induction, a BMP signaling gradient was established from the end producing BMP4 to the opposite end (Figures 6C, 6D), and the EB began to elongate progressively, reaching a length of about 0.5 mm after 5 days of development, with segments observable at this stage (Figures 6B, S6A). The elongation of EB arrested at day 6 (Figures 6A, 6B). Further assessment expression patterns of key markers in BMP4-induced EB revealed similarities to vertebrate caudal structures. We observed NMPs (TBXT^+^ and SOX2^+^) at day 4, and segmented somitic mesoderm (MEOX1^+^) at day5 in BMP4-induced EBs (Figures 6E, 6F). In the vertebrate embryo, MESP2(35) is expressed in the anterior PSM and newly formed somites, and the expression of TBX18(36) is restricted to the anterior halves of the somites. In the BMP4-induced human EBs, the region expressing MESP2 was broad and opposed to the ectodermal region expressing SOX2, suggesting the presence of the PSM (Figures 6G). The expression of TBX18 was evident, although not in a striped pattern, (Figure 6H), indicating that somites are formed.

**Figure 6.**
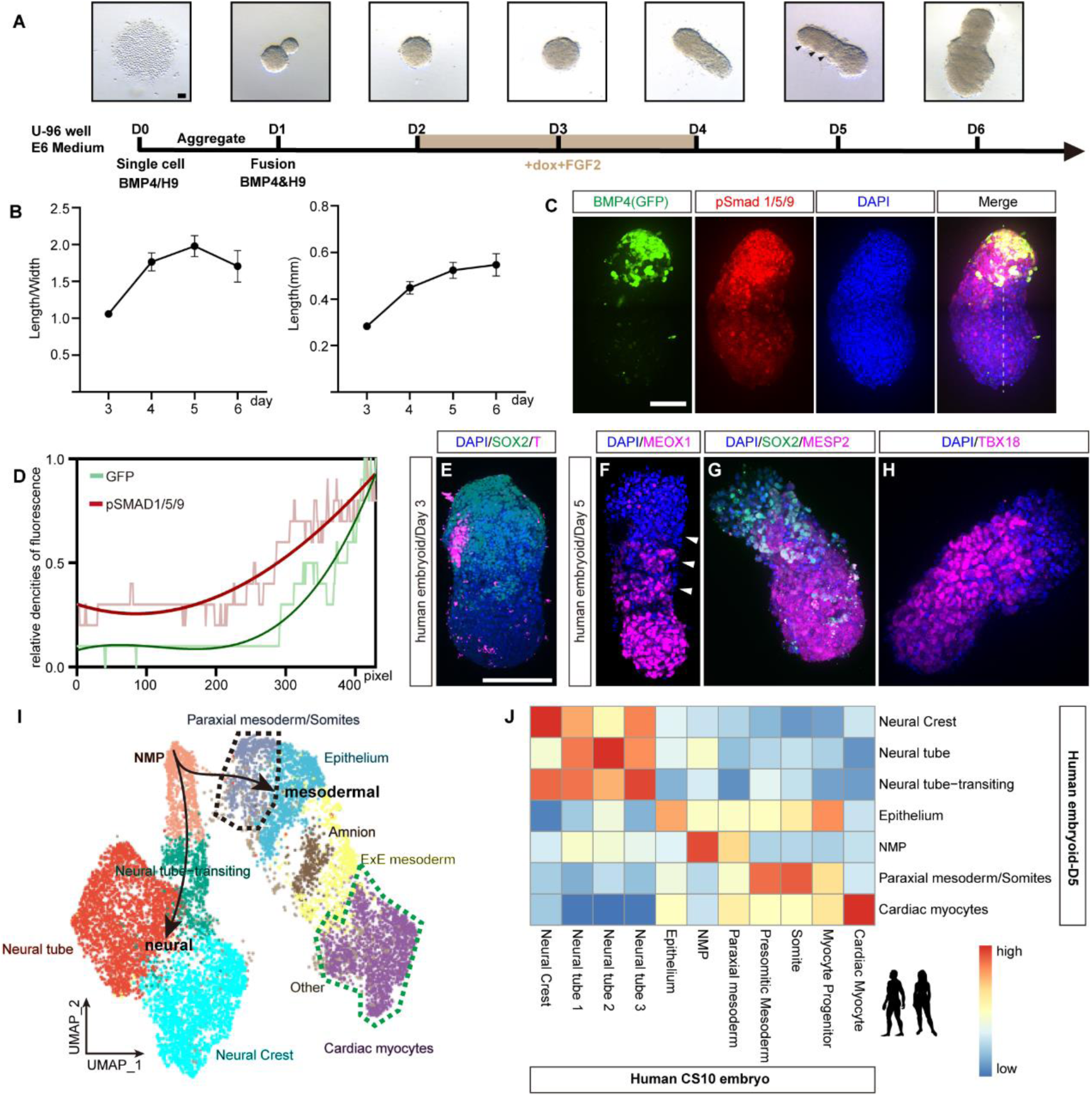
Construction of human caudal-like structure (CLS) with BMP4. (A) Representative images depicting the method used to induce human CLS with BMP4. (B) Quantification analysis of the length (right) and length/width ratio (left) of CLS. (C) Immunofluorescence staining of phosphorylated Smad1/5/9 and co-stained with DAPI in CLS. The BMP4 transgenic cells are labeled with GFP. (D) Quantification of pSmad1/5/9 and GFP signals along the dashed line shown in (C), normalized to their respective peak signals. (E-H) Immunofluorescence staining for T and SOX2 (E), MEOX1 (F), MESP2 and SOX2 (G), TBX18 and T (H) with DAPI co-stained in CLS. (I) UMAP plot showing the cell clusters in CLS at day5, with cells colored according to cell type. Black lines with arrows depict potential differentiation trajectories from NMP cells towards neural or mesodermal lineages. (J) Heatmap showing the Pearson correlation between gene expression profiles in human embryo and CLS. Scale bars:100 μm.

To further analyze the cellular and molecular features of BMP4-induced human EBs, we performed scRNA-seq analysis. At Day 5, we identified cell clusters such as NMPs, paraxial mesoderm/somites, neural tube and cardiac myocytes that resemble the caudal counterparts *in vivo* (Figures 6I, 6J). For example, the paraxial mesoderm/somites cell cluster in the EB displays high expressions of *MEOX1* and *CYP1B1*, and the cardiac myocytes cell cluster displays high expressions of *MYOCD* and *FN1,* similar to the situation in human embryo at Carnegie stages 10(37) (Figures S6B-S6E). At day 6, we also identified NMPs that expresses *SOX2* and *NKX1-2*, paraxial mesoderm/somites that expresses *MEOX1* and *CYP1B1* and cardiac myocytes that expresses *MYOCD* and *FN1,* similar to the situation in Monkey embryo at Carnegie stages 8-11(38) (Figures S7A-S7E). However, the overall paraxial mesoderm/somites cell fates were greatly diminished and transcriptionally resembled epithelium (Figures S7A, S7F), suggesting that somitogenesis was not normally progressed from day 5 to day 6.

Taken together, above observations suggest that BMP4-induced EBs display cellular and molecular features of human caudal structures. Further modifications on culture conditions may improve the somitogenesis progress in this model system.

## Discussion

In evolution and embryonic development, key regulators such as morphogens and transcriptional factors are repeatedly utilized across various locations and developmental stages(39, 40). It is well established that BMP4 is critical for dorsal-ventral patterning, tailbud development and organogenesis, with its functions conserved across a variety of species(41). Here, for the first time, we demonstrate that BMP4 alone is sufficient to induce caudal-like structures of a vertebrate embryo both *in vivo* and *in vitro*. In this study, we have demonstrated that *bmp4* overexpression at the animal pole of zebrafish robustly induces the formation of a secondary tail, and after explantation, the resulting structure exhibited anterior-posterior (A-P) patterned caudal tissues but lacked anterior neural ectoderm, axial mesoderm, and endoderm. Using single-cell RNA sequencing analysis, we showed that this model system not only displays molecular features characteristic of caudal cell fates but also parallels the developmental trajectories of its *in vivo* counterparts(42).

Notably, analysis of morphogenetic cell movements in this caudal-like structure revealed remarkable similarities to embryonic gastrulation and tail elongation processes observed *in vivo*: during gastrulation, involution was prominent, driving the precursors of paraxial mesoderm and lateral plate mesoderm (LPM) anteriorly, thereby establishing the A-P axis of the caudal-like structure (CLS). Intriguingly, a transient blastopore was observed at the posterior of the CLS—a feature not typically visible in zebrafish embryos due to the presence of the yolk cell(43). As dramatic elongation commenced, we observed several classical morphogenetic cell movements, including the irregular movement of TMP cells, their integration into the anterior PSM, and the migration of neural cells towards the TMP. It is noteworthy that cell proliferation and convergence & extension in the formed somites and PSM likely contribute to the elongation of the anterior part of the CLS, as similarly described *in vivo*(44, 45).

The concept of ‘organizer’ was first introduced by Hans Spemann and Hilde Mangold in 1924. It traditionally refers to a region at the dorsal blastopore of the embryo that can induce a secondary axis when transplanted to the ventral side of a host embryo. Subsequent studies identified additional embryonic regions with organizing properties, such as the tail organizer in zebrafish, which arises from the ventral blastoderm margin and can induce a secondary tail when grafted to the animal pole(4, 5). Xenografting of BMP4 treated human PSCs to zebrafish animal pole led to the formation of a secondary caudal structure, demonstrating the function of ‘caudal organizer’. This substantiates the feasibility of using BMP4 to construct human caudal-like structures. We successfully induced caudal cell fates, such as neural mesodermal progenitors (NMPs) and somites, in human PSC aggregates. Our results indicate that BMP4 is capable of inducing the caudal organizer, and this function is conserved among vertebrates. However, after 5 days of development, the elongation and segmentation processes ceased in this human CLS. Further modification of culture conditions or incorporation of additional signaling factors may be tested to better recapitulate human caudal development.

Overall, our work demonstrates that BMP4 can robustly induce the caudal organizer, a function that is highly conserved across different species. To validate this crucial role of BMP4, we employed a variety of experimental systems, including the *in vivo* overexpression of *bmp4* at the animal pole in zebrafish, zebrafish explant systems, xenografting of human pluripotent stem cells (PSCs) into zebrafish embryos, and 3D aggregates of human PSCs. We believe that by combining single-cell transcriptomics and live imaging techniques, the research platforms established in this study can not only be effectively used to study the role of morphogens and other key molecules in embryonic development but also to facilitate research such as cell-cell communications, morphogenetic cell movements, as well as the construction of embryoids and organoids *in vitro*.

## Methods

### Animals

Zebrafish wide type AB line and Tg(*actb2*:H2B-GFP), Tg(*aldh1a2*:H2B-RFP) transgenic lines are used in this study. Embryos were dechorionated by Pronase, raised in 0.3× Danieau’s buffer, kept at 28.5℃ and staged according to Kimmel et al(25). The Institutional Review Board of Zhejiang University and the Animal Ethics Committee of the School of Medicine, Zhejiang University, approved the experimental protocol. The protocol number is ZJU20220375.

### Cell lines

The WA09 human embryonic stem cells (hESCs, H9, WiCell) and H9 hESCs with doxycycline-inducible expression of BMP4 (iBMP4) were used in this research. The iBMP4 hESCs were generated as follows: The transposon plasmid containing a tetracycline (TRE) inducible BMP4 cassette (PB-Tre-BMP4) and constantly expressing GFP was constructed. To generate iBMP4 hESCs, the PB-Tre-BMP4 and PB-blue plasmids were co-transfected into H9 using the P3 Primary Cell 96-well Nucleofector Kit (Lonza) and the 4D Nucleofector×Unit (Lonza) according to the manufacturer’s instructions. Cells after electroporation were recovered in mTeSR-1 medium supplemented with ROCK inhibitor Y276932 (10 μM; Selleckchem). 24 hours later, these cells were cultured in fresh mTeSR-1 medium until reaching approximately 70% confluency and then were digested with Accutase (A6964, Sigma-Aldrich) at 37°C for 5-8 minutes to obtain single cell suspension. 500 cells were then seeded in six-well plates and cultivated until forming colonies.

### Injection of *bmp4* mRNA into zebrafish embryos and construction of Bmp4 explants

*bmp4*-injected zebrafish embryos were generated by injecting *bmp4* (8-10 pg) mRNA into one animal pole blastomere at the 128-cell stage. The injected embryos were raised at standard conditions until reaching desired stages. Bmp4 explants were generated by cutting off the animal pole region of *bmp4* injected embryos at 512-cell stage in a Petri dish coated with 2% agarose gel, using a syringe needle (27G SUNGSHIM MEDICAL CO., LTD). Then the explants were cultured in Dulbecco’s modified Eagle’s medium (DMEM/F-12 with 10 mM HEPES, 1× MEM with non-essential amino acids, 7 mM CaCl2, 1 mM sodium pyruvate, 50 μg/mL gentamycin, 100 μM 2-mercaptoethanol, 1× antibiotic-antimycotic, 10% serum replacement) and kept at 28.5℃.

### Construction of human caudal-like structures

Human H9 hESCs and iBMP4 hESCs were cultured on Matrigel-coated 12-well plates in mTeSR-1 medium. Cells were grown for 48 to 72 hours until reaching an approximate 70% confluence, when they are appropriate for the formation of embryoid body. The following induction process utilized a conditional medium (TeSR-E6 medium supplemented with 0.5% N2 and 0.5% B27 to support cell growth).

On day 0, both wild-type H9 cells and iBMP4 cells were dissociated using accutase and subsequently seeded into 96-well U-bottom low attachment plates. The seeding density was set at 1,300 cells per well for H9 cells and 400 cells per well for iBMP4 hESCs, each in 50 μL of aggregation medium. This medium was further fortified with 10 μM of the ROCK inhibitor Y27632.

After 24-hour incubation period, the cells formed spherical aggregates in their respective wells. Subsequently, one aggregate from the H9 well and one from the iBMP4 well were manually transferred into the same well with 80 μL of initial conditional medium to promote fusion. During days 2 and 4, the conditional medium was replaced with 80 μL induction medium (conditional medium supplemented with 75 ng/mL Doxycycline and 50 ng/mL FGF2) to induce BMP4 expression. Starting from day 4, the induction medium was replaced with 80 μL conditional medium for continued growth and development.

### Immunostaining

Samples were fixed by 4% paraformaldehyde at 4℃ overnight. Incubate the samples with 150 mM Tris-Hcl (PH 9.0) at 70℃ for 15 min to retrieve antigen. Treat samples with blocking buffer (10% FBS, 0.1% Triton X-100,1× PBS) for 2 h, then incubate overnight at 4℃ with antibody diluted in blocking buffer. Anti-pSmad1/5/9 antibody (13820, Cell Signaling Technologies, 1:800), anti-SOX2 antibody (AF2018, R&D, 1:50), anti-MEOX1 antibody (orb618083, Biorbyt, 1:50), anti-MESP2 antibody (orb669191, Biorbyt, 1:50) and anti-TBX18 antibody (MAB63371, R&D, 1:50) was used for the immunostaining experiments. Samples were washed 6 times with PBSTr (0.1% Triton X-100,1× PBS) to remove primary antibody. Treat samples with blocking buffer for 1 h, then with secondary antibody diluted in blocking buffer (1:500) until desired signal tensity is achieved. DAPI co-staining was achieved by adding 0.1 μg/mL DAPI with secondary antibody. Wash samples 6 times with PBSTr to remove secondary antibody.

### Live imaging

Samples were transferred to a glass-bottom culture dish (Cellvis, D35-10-0-N), properly oriented in one drop of 0.1% low melting agarose and covered with the culture medium. The samples were then photographed using OLYMPUS CSU-W1 confocal microscope every 3 min.

### Whole mount *in situ* hybridization (WISH)

Probes were labeled with digoxigenin-11-UTP (Roche Diagnostics), and the substrate was NBT/BCIP. Samples were fixed by 4% (w/v) paraformaldehyde in DEPC treated PBS at 4℃ overnight. Dehydrate the samples with 100% methanol at - 20℃ overnight. Rehydrate samples by washing with successive dilutions of Methanol. Hybridize the samples with hybridization mix containing 30-50 ng probe at 70℃ overnight. Gradually exchange the hybridization mix to 2× SSC and then incubate in 0.2× SSC at 70℃ for 2 times. Replace 0.2× SSC to PBST by successive dilutions of 0.2× SSC. Incubate samples in blocking buffer for 2 h and then in blocking buffer containing 1/5000 AP-anti-DIG antibody at 4℃ overnight. Remove the antibody by extensive washes of PBST. Incubate the samples in 1 mL staining solution (alkaline tris buffer, 225 μg/mL NBT, 175 μg/mL BCIP) until desired signal density was achieved. Block the labeling by washing with PBST (0.1% Tween-20,1× PBS) and stock the samples in glycerol.

### *In situ* hybridization chain reaction (HCR)

HCR staining was performed as described previously(46). Samples were fixed by 4% (w/v) paraformaldehyde in DEPC treated PBS at 4℃ overnight. Proteinase K treatment is recommended before incubation of probe sets. Samples were incubated with 2 pmol of each HCR probe sets in 100 μL probe hybridization buffer at 37℃ for 12-16 h. Wash with probe wash buffer at 37℃ for 4 times and with 5× SSCT at room temperature for 2 times to stop hybridization. Pre-amplify samples with amplification buffer at room temperature for 30 min, then incubate samples with 30 pmol snap-cooled hairpin h1 and h2 for 4 h at room temperature in a dark drawer. Stop amplification by washing with 5× SSCT (DAPI co-staining was achieved by adding 0.1 μg/mL DAPI) several times. Fluorescent hairpins, buffers and DNA probes were purchased from Molecular Technologies. Fluorescent signals were detected by OLYMPUS CSU-W1 confocal microscope.

### Xenograft assay

When iBMP4 cells were around 80% confluent, cells were dissociated using Accutase, washed in DMEM/F12, centrifuged at 180 g for 3 minutes and resuspended in mTeSR1 supplemented with 10 μM ROCK inhibitor Y27632. Cell numbers were determined using an automated cell counter and a calculated volume of single-cell suspension containing about 200,000 cells were added to a Matrigel pre-treated well of a 12-well plate. 24 hours after single-cell seeding, the culture medium was replaced with TeSR-E6 containing 1 μg/mL Doxycycline and 50 ng/mL recombinant Human FGF2, for a 24-hour treatment. Following treatment with Doxycycline, BMP4 treated cells were digested into single cells as described above and resuspended in 50 μL TeSR-E6. About 20-50 resuspended BMP4 treated cells were then transplanted to the animal pole of 4 hpf zebrafish embryos. The zebrafish embryos with induced secondary axis at 24 hpf were selected for imaging analysis and WISH.

### Processing of single-cell RNA-seq data

Raw sequencing reads were processed using the CellRanger(47) pipeline (10x Genomics, version 3.0.2) with default parameters. The resulting filtered expression matrix was utilized for downstream analysis, with the zebrafish reference genome GRCz11 or human reference genome GRCh38. Quality control and cell clustering were performed using the Seurat(48) package (version 4.0.3). Initially, a Seurat object was created with parameters min.cells = 3 and min.features = 200 by *CreateSeuratObject* function. Subsequently, the *subset* function was employed to remove cells based on appropriate parameters for each sample. Normalization of gene expression for each cell was performed using the *NormalizeData* function, and log-transformed expression counts were obtained. Highly variable genes were detected using the *FindVariableGenes* function, and the top 2000 genes with the highest variability were selected for downstream analysis. The data was scaled by running the *ScaleData* function. Linear dimensional reduction was performed by *RunPCA* function. Principal components were determined using the *JackStraw* and *ScoreJackStraw* functions. Clustering analysis was conducted using the *FindNeighbors* and *FindClusters* functions. The resolution parameter in the *FindClusters* function was set between 1 and 2. Differentially expressed markers for each cluster were identified using the *FindAllMarkers* function with default parameters. Finally, the cell clusters were annotated based on Zebrafish Information Network (bmp4 explant data, https://www.zfin.org/) or other published studies(38, 42, 49, 50).

### Integration of single-cell RNA-seq data

To integrate the Seurat objects of bmp4 explant of different timepoints or with wild-type embryonic dataset, each Seurat object was normalized by *NormalizeData* function. And then, the *FindVariableFeatures* function was used to detect the variable genes in each sample, the nfeatures was set to 2000. Next, select features that are repeatedly variable across all datasets that need to be integrated by running *SelectIntegrationFeatures* function. The identified features were used to run *ScaleData* and *RunPCA* for each object. Then, *FindIntegrationAnchors* function was used to determine the “anchors” of these objects. Finally, the integrated dataset was obtained by running *IntegrateData* function. After that, the default assay was set to ‘integrated’, and the standard workflow for visualization and clustering was executed, including the following steps: *ScaleData*, *RunPCA*, *RunUMAP*, *FindNeighbors*, and *FindClusters*. The identified cell clusters in integrated Seurat object were visualized on UMAP(51) plot.

### Construction of single-cell trajectory for Bmp4 explants

Three different methods were employed to conduct single-cell RNA-seq trajectory analysis. Firstly, URD(42) was employed to reconstruct the developmental trajectories of Bmp4 explant cells, spanning the time range from 6 to 24 hpf. The analysis was conducted following the instruction of previous study(42). In this analysis, early time point cells (6 hpf) were designated as the root, while late time point cells (24 hpf) were assigned as the tips of the trajectory. The tip clusters were annotated based on the differentially expressed markers within each cluster. To construct the branching tree, we utilized the *buildTree* function in URD with the following parameters: save.all.breakpoint.info = T, divergence.method = “preference”, cells.per.pseudotime.bin = 50, bins.per.pseudotime.window = 5, and p.thresh = 0.01. The branchpoint preference plot for TMP and PSM was generated using the *branchpointPreferenceLayout* function from URD. Subsequently, stage, pseudotime information, and gene expression were visualized on the branchpoint using the *plotBranchpoint* function. For identifying the deferential expression (DE) genes along the trajectories, we utilized the *geneCascadeProcess* function with a moving window average (moving.window = 5, cells.per.window = 15). This smoothing process enabled us to obtain a more refined representation of gene expression dynamics. The smoothed expression of each gene was fitted using an impulse model.

Next, NMP cells, as well as neural derivatives and mesodermal derivatives, were selected for cell differentiation analysis using monocle3(52) with default parameters. This analysis aimed to demonstrate the bipotent differentiation trajectory of NMP cells. Similarly, cells from the TMP, PSM, and somite populations were selected for analysis using monocle3. This analysis allowed for the identification of different gene modules along the TMP-PSM trajectory and the PSM-somite trajectory. *graph_test* function was used to identify DE genes along the trajectories and *find_gene_modules* function was employed to detect related gene modules that regulate cell differentiation along specific trajectory.

Lastly, a K-Nearest Neighbors (KNN) algorithm-based method was employed to construct single-cell developmental trajectory. The scRNA-seq data from adjacent time points was integrated using Harmony(53) with default settings. Then the KNN algorithm was used to select the 10 nearest neighbors in the earlier time point for each spot of the latter time point. The most frequent clusters of the 10 nearest spots were regarded as the source states that the latter spots would develop from. If two or more clusters had the same frequencies, the cluster of the nearest spot was used. The procedure was repeated on each pair of adjacent time points from 6 hpf to 24 hpf, and a connection was only reserved when there were more than 20% cells of the same cluster in the latter time point developing from a cluster in the earlier time point.

### Comparison with *in vivo* datasets

The scRNA-seq datasets of Human CS10 embryos(50) and Monkey CS8-CS11 embryos(38) were used as references for comparison. The gene expression of each cluster in the human embryoids and the reference datasets was aggregated into pseudo-bulk. The marker genes of each cluster were selected, and a correlation test was performed using the marker gene expression in the pseudo bulk to assess the cell type similarities between the human embryoids and the references. The correlations of cell types between the human embryoids and the references were calculated as Pearson correlation coefficients and visualized using a heatmap.

### Gene set enrichment analysis

The genes from gene modules along the TMP-PSM trajectory or PSM-somite trajectory were used to perform gene set enrichment analysis. The gene list was uploaded to the Metascape(54) website (http://metascape.org), and the results were downloaded following the execution of Express Analysis.

### Tracking cell movements in Bmp4 explants

Imaris software (version 9.7) was utilized to and track the cell movements in bmp4 explants, or the embryos injected with *bmp4* mRNA. The Spot plugin from Imaris was employed to label the cells and analyze cell trajectories. Spot detection diameter was set at 6–15 μm (Figure S5C-5D, 15μm; S5E-5F and 4B, 13.5 μm; S3G-3H, 6.5 μm). Subsequently, candidate spots were manually selected, filtering out spots that met the appropriate quality threshold, for further analysis. Autoregressive Motion algorithm was employed to track the movement of each cell, with Max Gap Site set to 3. The movement dataset, consisting of cell coordinates at all time points, were extracted from Imaris and analyzed in R. It was observed that all cells exhibited migration in the same direction. This migration pattern, referred to as the observed trajectory (OT), was primarily attributed to the elongation of the entire explant. To analyze the cell trajectory specifically within the Bmp4 explant, termed the migration trajectory (MT), the elongation trajectory (ET) was first derived from the OT by studying the movements of EVL cells (designated as anchor cells) that were presumed to remain stationary within the bmp explant. Three anchor cells (EVL cells) were selected, and their central coordinate trajectory was defined as ET. Finally, the MT of the selected cells was obtained by subtracting the ET from the OT. The trajectories were visualized by R package plot3D (https://CRAN.R-project.org/package=plot3D).

## Data availability

The data generated in this study is deposited in the Gene Expression Omnibus (GEO). The single-cell RNA sequencing data is under accession number GSE270989 (https://www.ncbi.nlm.nih.gov/geo/query/acc.cgi?acc=GSE270989).

## Code availability

Any custom code and data are available from the authors upon request. All analyses are based on previously published code and software.

## Supplemental information

Supplementary Figures 1-7, Supplementary Table 1-4, Supplementary movies 1-11

## Acknowledgements

We thank Dr. Bernard Thisse (1959-2021) and Dr. Christine Thisse at University of Virginia for their helpful comments and suggestions. We also thank Dr. Min-Xin Guan, Dr. Xiao-Hang Yang, Dr. Hong-Qing Liang and the members of Laboratory of Development and Organogenesis (LDO) at Zhejiang University for helpful suggestions and discussions. This work was supported by grants from the National Scientific Foundation of China (31900576, 32300677, 32300688) and the Chinese National Key Research and Development Project (2022YFA1103102, 2019YFA0802402).

## Declaration of interests

The authors declare no competing interests.

## Author Contributions

PF X, YYX, TC and YH designed the research; YYX, YH, TC, TL, YD, ZXJ, YMT and XL performed research; TC and YD analyzed bulk RNA-seq and single cell RNA-seq data; YYX, PFX and TC wrote the manuscript; JM and JFJ revised the manuscript; all authors reviewed and approved the manuscript.

